# Positive sexual imprinting for human eye color

**DOI:** 10.1101/135244

**Authors:** Lisa M. DeBruine, Benedict C. Jones, Anthony C. Little

## Abstract

Human romantic partners tend to have similar physical traits^1^, but the mechanisms causing this homogamy are controversial. One potential explanation is direct matching to own characteristics^2,3^. Alternatively, studies showing similarity between parent and partner^4,5^ support positive sexual imprinting^6,7^, where individuals are more likely to choose mates with the physical characteristics of their other-sex parent. This interpretation has been strongly criticized because the same pattern could also be caused by sex-linked heritable preferences^3^, where similarity in appearance between an individual’s partner and their other-sex parent is caused by similarity in preferences between the individual and their same-sex parent. The relationships among own, parents’ and same-sex partner’s eye color provide an elegant test of these hypotheses, which each postulate a different best predictor of partner’s eye color. While the matching hypothesis predicts this will be own eye color, the sex-linked heritable preference hypothesis predicts this will be the other-sex parent’s eye color and the positive sexual imprinting hypothesis predicts this will be the partner-sex parent’s eye color. Here we show that partner eye color was best predicted by the partner-sex parent’s eye color. Our results provide clear evidence against matching and sex-linked heritable preference hypotheses, and support the positive sexual imprinting hypothesis of the relationship between own and partner’s eye color.

---

Humans exhibit homogamy in mate choice and preferences across a range of personality and physical characteristics^1^, including facial appearance^2,8,9^, hair color^4^, and eye color^4,5^. Explanations for this phenomenon fall into three main categories: those that emphasize matching to own characteristics^1^, those that emphasize learning from parental models^4,6,9^, and those that emphasize the transmission of preferences from parents^3^.

Two main phenomena relate to directly matching own characteristics: self-referential phenotype matching and assortative mating. Self-referential phenotype matching involves sampling one’s own phenotype and using it to inform social behavior either positively or negatively. For example, cross-fostered ground squirrels raised apart from genetic family members recognize their unfamiliar genetic siblings, but not foster siblings, after hibernation^10^, presumably because of similar phenotypic characteristics such as odor. Homogamy can also happen in the absence of a preference for self-similar traits^11^. For example, if a characteristic is generally preferred, regardless of own phenotype, the most desirable mates are more likely to both have the characteristic and to be able to attract a mate with the characteristic, leading to assortative mating even in the absence of assortative preferences.

Positive sexual imprinting can lead to resemblance between mates insofar as offspring are similar to their parents. Acquiring mate preferences from a parental model may facilitate speciation^12^. For example, subspecies of zebra finch have been shown to avoid hybridization by preferring the songs and plumage of their own species^13^. Additionally, cross-fostered females mated with males who were similar to their foster-fathers^13^. Parental imprinting can be sex-specific, as elegantly demonstrated by a study of cross-fostered sheep and goats; cross-species fostering had a strong and irreversible effect on male mate choice, while it had a weaker and reversible effect on female mate choice^14^. This sex difference in ungulate species is likely to be a result of only experiencing maternal care. In a bi-parental species such as humans, parental-imprinting-like effects in both men and women have been shown to be relatively specific to the other-sex parent. For example, other-sex parental hair and eye color are significantly better predictors of partner hair and eye color than are same-sex parental hair and eye color^4^. Additionally, studies have found that, for men, their wives’ facial appearance is best predicted by their mothers’ facial appearance^8^, while for women, their husbands’ facial appearance is best predicted by their fathers’ facial appearance^9^.

The evidence for positive sexual imprinting in humans has been strongly criticized because sex-linked heritable preferences can also lead to similarity between one’s mate and other-sex parent^3^. While twin studies show evidence for substantial heritability of preferences for some traits such as height, hair color, and physical attractiveness^15,16^, the heritability of realized mate choice (as measured from actual partners) is very low^1^. However, this hypothesis is difficult to reconcile with consistent findings that the parent-child relationship affects the strength of preferences for faces that resemble one’s other-sex parent^17^ and the strength of resemblance between that parent and one’s spouse^8,9^, as well as husband-father resemblance being observed even among adopted women who were not genetically related to their fathers^9^.

We tested these three hypotheses by assessing participants’ own, parents’ and partner’s eye color. While the matching hypothesis predicts that the best predictor of partner’s eye color will be own eye color, the sex-linked heritable preference hypothesis predicts this will be the other-sex parent’s eye color and the positive sexual imprinting hypothesis predicts this will be the partner-sex parent’s eye color. While these latter two predictions lead to the same pattern of results for heterosexual couples, here we also test same-sex couples to distinguish the sex-linked heritable preference hypothesis from the positive sexual imprinting hypothesis.

**Figure 1.**
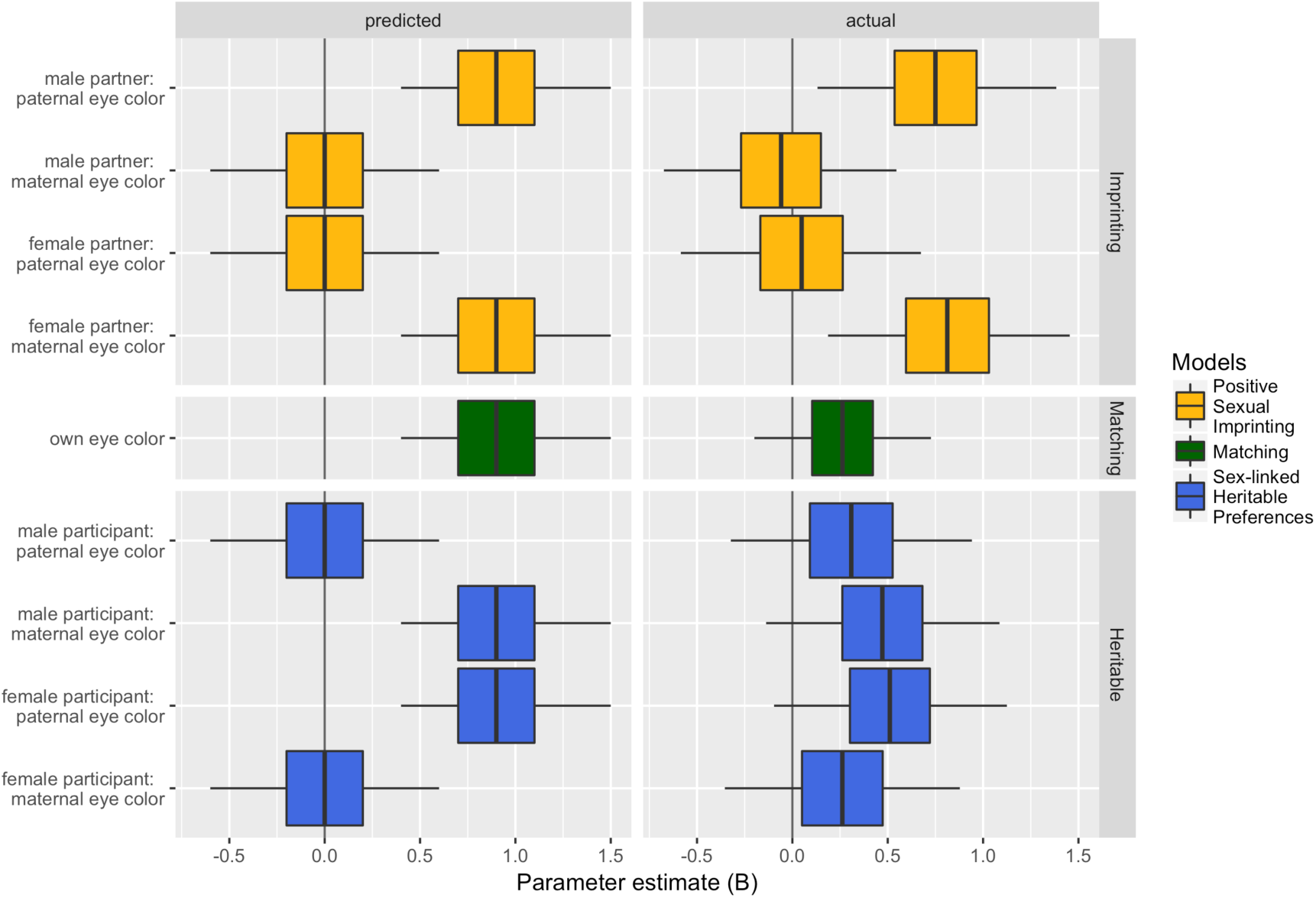
Model summaries. Predicted and actual parameter estimates for all 3 models. Boxes show the 50% CI and whiskers show the 95% CI. Predicted models are estimated from simulated data with N = 300 and effect sizes from previous research^4^.

We used binomial logistic regression to test three models representing the predictions of our three hypotheses and Akaike Information Criterion^18^ to compare the models (see Table 1). The best model supported the positive sexual imprinting hypothesis, where maternal eye color significantly predicted the eye color of female partners (B = 0.812, S.E. = 0.322, z = 2.52, p = 0.012), but not male partners (B = - 0.059, S.E. = 0.31, z = -0.19, p = 0.849) and paternal eye color significantly predicted the eye color of male partners (B = 0.75, S.E. = 0.318, z = 2.355, p = 0.019), but not female partners (B = 0.049, S.E. = 0.32, z = 0.152, p = 0.879). This model was supported 7.455 times more strongly than the model corresponding to the matching hypothesis, where own eye color did not significantly predict partner eye color (B = 0.263, S.E. = 0.236, z = 1.113, p = 0.266). The imprinting model was also supported 9.655 times more strongly than the model corresponding to the sex-linked heritable preference hypothesis, where maternal eye color did not predict the eye color of men’s partners (B = 0.262, S.E. = 0.313, z = 0.836, p = 0.403) more than women’s partners (B = 0.472, S.E. = 0.311, z = 1.515, p = 0.13) and paternal eye color did not predict the eye color of women’s partners (B = 0.309, S.E. = 0.321, z = 0.96, p = 0.337) more than men’s partners (B = 0.511, S.E. = 0.311, z = 1.645, p = 0.1). Indeed, the opposite of the predicted pattern was found, although none of the individual predictors were significantly related to partner eye color.

**Table 1.** Model comparison for binomial models.

| Model | Parameters | $K$ | $\Delta_i$ | $w_i$ | Evidence Ratio w/<br>imprinting |
| --- | --- | --- | --- | --- | --- |
| imprinting | maternal:partnersex +<br>paternal:partnersex | 5 | 0 | 0.696 | 1.00 |
| null | (Intercept) | 1 | 3.232 | 0.138 | 5.033 |
| matching | own | 2 | 4.018 | 0.093 | 7.455 |
| heritable | maternal:ownsex +<br>paternal:ownsex | 5 | 4.535 | 0.072 | 9.655 |

$K$ = the number of parameters in the model; $\Delta_i$ = change in $AIC_c$ from the best model; $w_i$ = Akaike weight (i.e., the relative likelihood of the model).

We also repeated these analyses using an 8-level ordinal coding of eye color. The model supporting the positive sexual imprinting hypothesis remained the best model (see Table 2).

**Table 2.** Model comparison for ordinal models.

| Model | Parameters | $K$ | $\Delta_i$ | $w_i$ | Evidence Ratio w/<br>imprinting |
| --- | --- | --- | --- | --- | --- |
| imprinting | maternal:partnersex +<br>paternal:partnersex | 11 | 0 | 0.818 | 1.00 |
| matching | own | 8 | 4.574 | 0.083 | 9.846 |
| null | (Intercept) | 7 | 5.491 | 0.053 | 15.573 |
| heritable | maternal:ownsex +<br>paternal:ownsex | 11 | 5.743 | 0.046 | 17.666 |

Our data provide clear evidence against the sex-linked heritable preferences hypothesis^3^, which predicts a relationship between partner’s eye color and other-sex parent’s eye color. In fact, our data show a non-significant trend in the opposite direction, with partner’s eye color being better predicted by same-sex parent’s eye color. Our data also give little support to the matching hypothesis, as own eye color was positively but not significantly related to partner’s eye color. Our data give clearest support to the positive sexual imprinting hypothesis, where we found that partner’s eye color was predicted by maternal eye color for people with female partners and by paternal eye color for people with male partners.

## METHODS

### Participants

We report data from women (n = 150, mean age = 22.9 years, SD = 7.91) and men (n = 150, mean age = 28.3 years, SD = 10.46) who completed an online questionnaire about eye color. Data were limited to individuals who reported own, maternal, paternal and partner eye color, as well as own and partner’s sex, and who reported on a biologically related mother and father (e.g., adoptees were excluded). Because of differences in the relative proportions of same-sex and different-sex relationships, data were collected continuously until we reached 75 individuals meeting the criteria above in each of four categories: women reporting on a female partner, women reporting on a male partner, men reporting on a female partner, and men reporting on a male partner.

Data collection was completed through a custom-built website (faceresearch.org) that participants typically access via social bookmarking links (e.g., stumbleupon.com). Participants were not compensated for their participation. The study complies with ethical regulations of the British Psychological Society, informed consent was obtained from all participants, and the study’s protocol was approved by the University of Glasgow’s psychology ethics board.

### Eye color

Following previous work on imprinting and eye color^4^, participants were asked to choose eye colors from the following list: black, dark brown, light brown, hazel, green, blue green, blue, and grey. For binomial analyses, the first three categories were recoded as “dark” and the last five categories were recoded as “light”, following previous work^4^.

### Statistical tests

Statistical tests were conducted using R 3.3.2^19^. To investigate which factors best predicted partner eye color, we used a set of a priori models based on the three hypotheses of matching, imprinting, and sex-linked heritable preferences. We conducted GLMs with binomial error structure. The dependent variable in each case was partner eye color. The null model included only an intercept. The additional predictor for the matching hypothesis was own eye color. The additional predictors for the parental imprinting hypothesis were maternal and paternal eye color, both interacting with partner sex. The additional predictors for the sex-linked heritable preferences hypothesis were maternal and paternal eye color, both interacting with participant sex. We compared these four models using the AICcmodavg package^20^. We also repeated these analyses using an ordinal coding of eye color (black = 1, dark brown = 2, light brown = 3, hazel = 4, green = 5, blue green = 6, blue = 7, and grey = 8) and GLM with Poisson error structure.

**Figure 2.**
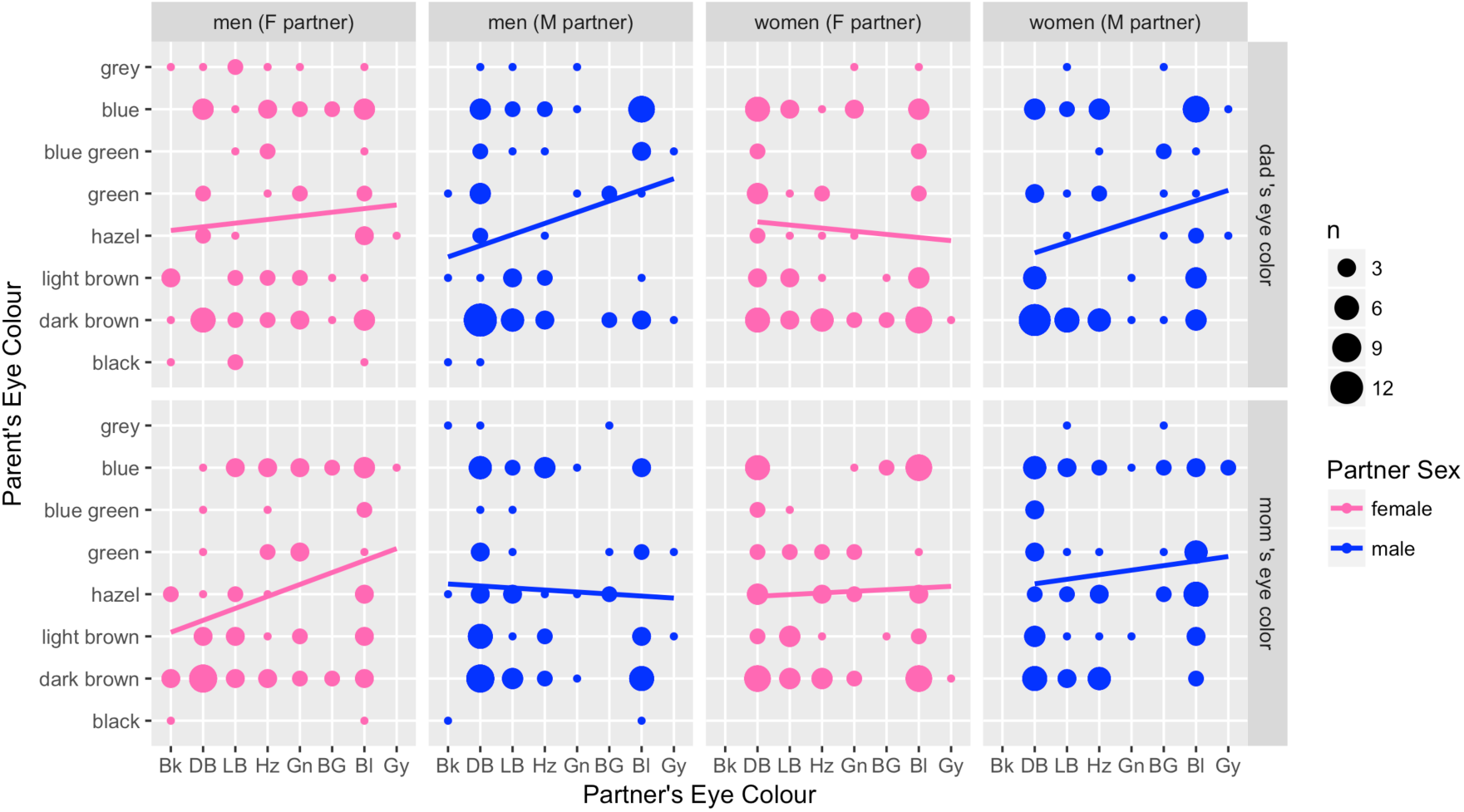
Data for ordinal coding of eye color. Blue represents data for male partners, pink represents data for female partners, and dot size represents number of participants (total N = 300).

In addition to the main analyses reported above, we also followed the exact backwards conditional binomial logistic regression analyses from previous work^4^, investigating how maternal, paternal and own eye color predicted partner eye color in each group of participants separately. We replicated the findings that women’s male partners’ eye color was best predicted by paternal eye color (B = 1.052, S.E. = 0.481, Z = 2.188, p = 0.029), while men’s female partners’ eye color was best predicted by maternal eye color (B = 1.072, S.E. = 0.486, Z = 2.207, p = 0.027). We also found that that men’s male partners’ eye color was best predicted by paternal eye color (B = 0.847, S.E. = 0.478, Z = 1.774, p = 0.076). While women’s female partners’ eye color was best predicted (negatively) by own eye color (B = -0.827, S.E. = 0.489, Z = 2.188, p = 0.091), maternal eye color was the only positive predictor in the first step of this analysis (B = 0.694, S.E. = 0.537, Z = 1.292, p = 0.196).

## Data availability

All data and reproducible analysis code are available at osf.io/he4ty^21^.

## Acknowedgements

This research was supported by ERC grant #647910 KINSHIP to L.M.D.

## Author Contributions

L.M.D. designed the study, conducted the analyses, and drafted the paper. B.C.J. consulted on theory and revised the paper. A.C.L. consulted on theory and analyses, and revised the paper.

